# D^3^M: Detection of differential distributions of methylation levels

**DOI:** 10.1101/023879

**Authors:** Yusuke Matsui, Masahiro Mizuta, Satoru Miyano, Teppei Shimamura

**Affiliations:** Nagoya University Graduate School of Medicine, 466-8550, Nagoya, Japan; Information Initiative Center, Hokkaido University, 060-0811, Sapporo, Japan; Institute of Medical Science, The University of Tokyo, Tokyo, 108-8639, Japan

## Abstract

**Motivation:** DNA methylation is an important epigenetic modification related to a variety of diseases including cancers. We focus on the methylation data from Illumina’s Infinium HumanMethylation450 BeadChip. One of the key issues of methylation analysis is to detect the differential methylation sites between case and control groups. Previous approaches describe data with simple summary statistics and kernel function, and then use statistical tests to determine the difference. However, a summary statistics-based approach cannot capture complicated underlying structure, and a kernel functions-based approach lacks interpretability of results.

**Results:** We propose a novel method D^3^M, for detection of differential distribution of methylation, based on distribution-valued data. Our method can detect high-order moments, such as shapes of underlying distributions in methylation profiles, based on the Wasserstein metric. We test the significance of the difference between case and control groups and provide an interpretable summary of the results. The simulation results show that the proposed method achieves promising accuracy and shows favorable results compared with previous methods. Glioblastoma multiforme and lower grade glioma data from The Cancer Genome Atlas show that our method supports recent biological advances and suggests new insights.

**Availability:** R implemented code is freely available from

https://github.com/ymatts/D3M/

https://cran.r-project.org/package=D3M.

## 1 INTRODUCTION

DNA methylation is an epigenetic chemical alternation in which a methyl group is attached to a 5-carbon of a cytosine (C) base. It is closely related to gene expression, silencing, and genomic imprinting, including oncogenesis. Typically, methylation is explained as occurring in cytosine-phosphate-guanine (CpG) sites. The methylation of promoter regions, in particular, silences cancer suppressor genes (Baylin, 2005; Kulis and Esteller, 2010).

We focus on the methylation data from Illumina’s Infinium HumanMethylation450 BeadChip. One of the key issues for methylation analysis is to detect differentially methylated sites, *i.e.,* a significant difference in methylation levels between the case and the control groups at a site. When comparing groups, we often summarize (or aggregate) data in summary statistics, such as mean and variance, and then investigate the difference between the groups. For example, IMA (Wang, *et al.*, 2012) detects the differentially methylated sites by T-test, Empirical Bayes (EB) method or by Mann–Whitney–Wilcox (MWW) test. DiffVar (Phipson, *et al.* 2014) detects the sites by testing significant difference of variance. Other nonparametric approaches exist, such as the Kolmogorov–Smirnov test (KS) or kernel-based approaches, such as maximum mean discrepancy (MMD) (Gretton, *et al.*, 2012). In particular, since KS and MMD consider the underlying distribution structure, they are better suited for use with complicated distributions than methods based on summary statistics.

These approaches are effective in detecting typical differential methylation sites, but are insufficient from some perspectives; underlying distributions are complicated by being skewed, heavy-tailed, and multimodal. In particular, since cancer cells include heterogeneities, measurements of the methylation levels potentially include complex distribution shapes. This observation indicates that we need to consider the underlying structure. The disadvantage of KS and MMD is infeasible interpretability of results because they measure the maximum and kernel distances of distributions, respectively, which are difficult to interpret corresponding to the actual difference of underlying distributions.

We develop a method to detect differential methylation sites with distribution-valued data (Irpino and Verde, 2014). Distribution-valued data are an example of symbolic data analysis (Diday, 1989). This framework can treat complex data such as functional (Ramsey and Silverman 2005), tree (Wang and Marron, 2007), set, interval, and histogram values (Bock and Diday, 2000). The proposed method describes case and control groups using distribution values. We measure the differences between distributions using the Wasserstein metric. We detect the differential methylation sites using a statistical test of significant differences of distribution functions.

## 2 METHODS

Our method is aimed at a distribution-based comparison of methylation levels in two groups, through site-by-site resolution. We construct distribution functions representing the two groups at each site. Next, we compare the groups using a dissimilarity measure and test statistical significance through site-by-site resampling. We adopt an *L*_2_-Wasserstein metric (Ruesehen, 2011) as a dissimilarity measure, a distribution function-based measure of statistical distance. The advantage of this distance is the interpretability of results because the distance can be decomposed into three components, *i.e.,* mean, variance, and distribution shape. This fact leads to visualization of results using a Q-Q plot to interpret the detected distribution difference including hypo- or hyper- methylation status. In the following equation, we assume that input data is a beta value that is the ratio between the methylated probe intensity and the overall intensity (Du, *et al.*, 2010). The definition of *i*-th site beta value in Illumina methylation assay is as follows:

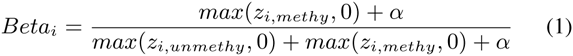

where *z_i,methy_* and *z_i,unmethy_* are the intensity measured by *i*-th methylated and unmethylated probes, respectively and α is constant.

### 2.1 Construction of objects

*X(s_i_)* and *Y*(*s_i_*) *(i* = 1, 2,…, *S*) represent the beta values in a case group (e.g., cancer subjects) and a control group (e.g., normal subjects) at a CpG site *s_i_*. We represent the data as distribution values by

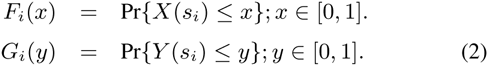

In practice, let the beta value observations be *x_j_*(*s_i_*); *j* = 1, 2,…, *n* and *y_j_* (*s_i_*); *j* = 1, 2,…, *m* following *F_i_*(*x*) and *G_i_*(*y*), respectively, where *n* and *m* are the respective numbers of observations at *s_i_*. From the data, we construct the empirical distribution functions;

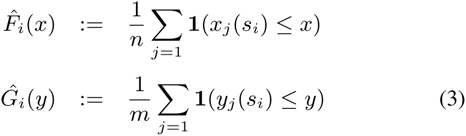

where

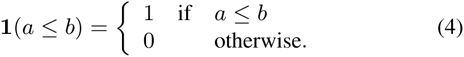

### 2.2 Dissimilarity measure for distributions

The Wasserstein metric is defined by the following:

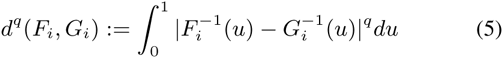

where 1 ≤ *q* ≤ 2 and 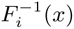 and 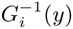 indicate quantile functions.

In particular, in the case of *q* = 2, the metric can be decomposed into three components that describe the distribution characteristics, *i.e*., mean, variance, and shape (Irpino and Verde, 2014):

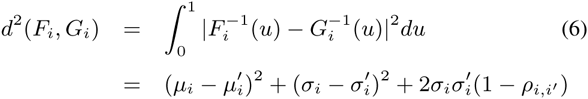

where *μ_i_* and 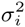 (respectively, 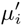 and 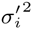) are mean and variance of *F_i_*(*x*) (respectively, *G_i_*(*y*)), and *ρ_i_* is the correlation index of the points in the Q-Q plot of *F_i_* and *G_i_*.

The empirical estimator of the Wasserstein metric is given by

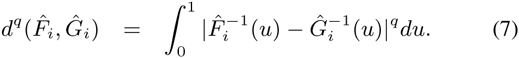

Technically, we use quantiles to compute the approximation of the (7) for reducing computational costs. Let (*Q_i_*,_1_, *Q_i,_*_2_,…, *Q_i_*,*_K_*) and 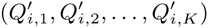 be *k*-quantiles of *F_i_*(*x*) and *G_i_*(*y*). We calculate 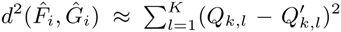 in the case of *q* = 2, instead of evaluating the integral in (7). Here we simply write 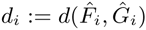.

### 2.3 Detection of differential methylation sites

We use the metric to investigate whether two distributions are significantly different. We pose statistical hypotheses as follows.

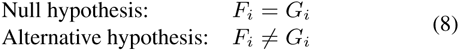

We use resampling to construct a null distribution. From the null hypothesis (8), we jointly permute the observations (*x*_1_(*s_i_*), *x*_2_(*s_i_*),…, *x_n_*(*s_i_*)) and (*y*_1_(*s_i_*), *y*_2_(*s_i_*), …, *y_m_*(*s_i_*)) to obtain the new distribution functions 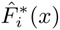 and 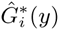. Next, we obtain the new distance 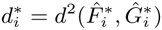 according to (7).

Let 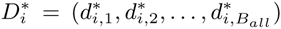 be all possible distances for the permutation process. Then *p*-value is

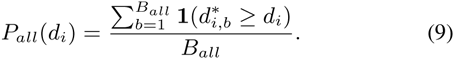

Approximation of (9) uses the subset of 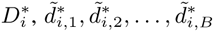 where B ≤ B*_all_*:

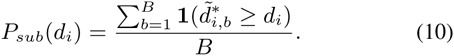

In the simulation of section 3 and data analysis in section 4, we set B = 10000.

The number of permutations *B* is closely related to the accuracy of the *p*-value. However, resolution of *P_sub_* is limited to 1/*B*, if we need the very small *p*-values. One solution is to perform a large number of permutations, but it is computationally expensive. A semi-parametric estimation of *p*-value is proposed by Knijnenburg *et al.* (2009) to obtain more accurate *p*-values.

We use an exponential distribution to estimate the distribution tail as follows,

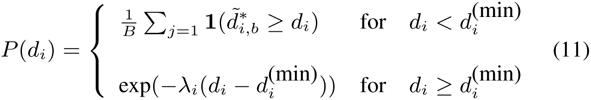

where *λ_i_* is a scale parameter and 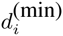 is a threshold that we set to 99^th^ percentile of null distributions. We estimate *λ_i_* using data above the threshold. Technically, we perform the semi-parametric estimation only if *P_sub_*(*d_i_*) reaches to zero.

### 2.4 Graphical representation of results

The graphical interpretation of the statistical test result is important. One approach is to plot all the distribution (density) functions of candidate sites, but this is infeasible for hundreds of sites. We use a Q-Q plot with two distributions. It enables us to visualize many pairs of distributions at a time, with the directions being easy to interpret. In the actual example shown in section 4, we plotted 1,000 pairs of differentially methylated distributions (Figure 3 (B)). We can see the hyper-methylation with the most significant 1,000 sites (blue lines in Figure 3 (B)).

## 3 SIMULATION

### 3.1 Simulation setting

We evaluated the proposed method with simulated datasets with focus on single cytosine level in the case and control group. Our simulation is intended for the detection of differential methylation sites when there is cancer heterogeneity. Here, the cancer heterogeneity is described by the multiple modes of distributions. We conduct a statistical test for *H*_0_ *: F_i_ = G_i_ ↔ H*_1_ *: F_i_ ≠ G_i_* under significance levels 5%, and we compare the results to those of the other methods, *i.e.,* DiffVar, MMD, KS, MWW and EB. We used several packages; `missMethyl` (Phipson, *et al.*, 2014), `limma` (Ritchie, *et al.*, 2015) written in R and `mmd` (Gretton *et al.*, 2006) (http://www.kyb.tuebingen.mpg.de/bs/people/arthur/code/mmd.zip) written in MATLAB. The setting of `missMethyl` is default and `mmd` with options `alpha = 0.5` and `MMD_METHOD = ‘approxmoments’`.

We describe the outline of the simulation as follows. The details are described in supplemental file S1 and R codes are described in S2. The data are generated by using two types of distributions. The control and case groups are represented by normal and normal mixture distributions, respectively. In each case (case 1–case 8), there are 80 samples; 40 samples for case and control groups, respectively. We performed the statistical test with each a method for 50 times in every case (*i.e.,* case 1–case 8) and evaluate averages of a type I error and power. Besides we repeat this process for 100 times to assess the variances of the averages.

### 3.2 Simulation results

The results are shown in Table 2. In the first case, it is shown that error rates of D^3^M, MMD, DiffVar, KS, EB, and MWW are close to the significance levels, which indicates that they effectively control type I errors.

**Table 1.** Simulation models of eight cases

| | $F_i = F_2$ | $F_1 \neq F_2$ | | | | | | |
| --- | --- | --- | --- | --- | --- | --- | --- | --- |
|  | case 1 | case 2 | case 3 | case 4 | case 5 | case 6 | case 7 | case 8 |
| $\mu_1 = \mu_2$ | T | T | T | F | T | F | F | F |
| $\sigma_1 = \sigma_2$ | T | T | F | T | F | T | F | F |
| $s_1 = s_2$ | T | F | T | T | F | F | T | F |

**Table 2.**
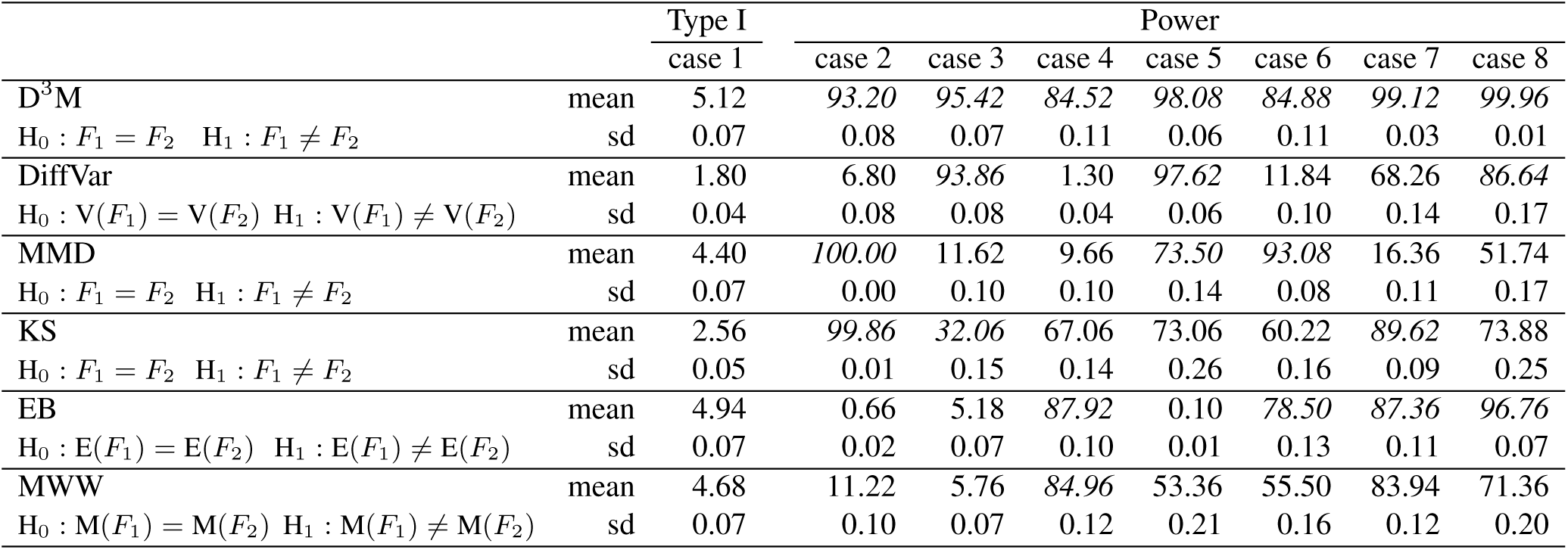
Hypothesis testing in each case (hypotheses are described under the method names). We focus on a probe and generate beta values, 40 samples each in a case and a control group respectively. We apply six methods to this data and evaluate the p-value at the significance level, *i.e.,* 5%. This process is repeated 50 times and returns the averages of type I error (%) and power (%). We represent the power in italics within top three. Furthermore, we also evaluate the standard deviations of the averages; we repeated the process of the evaluating average of type I error and power a 100 times and then obtained the standard deviations of the averages.

|  |  | Type I | Power |  |  |  |  |  |  |
| --- | --- | --- | --- | --- | --- | --- | --- | --- | --- |
|  |  | case 1 | case 2 | case 3 | case 4 | case 5 | case 6 | case 7 | case 8 |
| D<sup>3</sup>M | mean | 5.12 | 93.20 | 95.42 | 84.52 | 98.08 | 84.88 | 99.12 | 99.96 |
| H<sub>0</sub> : F<sub>1</sub> = F<sub>2</sub> H<sub>1</sub> : F<sub>1</sub> ≠ F<sub>2</sub> | sd | 0.07 | 0.08 | 0.07 | 0.11 | 0.06 | 0.11 | 0.03 | 0.01 |
| DiffVar | mean | 1.80 | 6.80 | 93.86 | 1.30 | 97.62 | 11.84 | 68.26 | 86.64 |
| H<sub>0</sub> : V(F<sub>1</sub>) = V(F<sub>2</sub>) H<sub>1</sub> : V(F<sub>1</sub>) ≠ V(F<sub>2</sub>) | sd | 0.04 | 0.08 | 0.08 | 0.04 | 0.06 | 0.10 | 0.14 | 0.17 |
| MMD | mean | 4.40 | 100.00 | 11.62 | 9.66 | 73.50 | 93.08 | 16.36 | 51.74 |
| H<sub>0</sub> : F<sub>1</sub> = F<sub>2</sub> H<sub>1</sub> : F<sub>1</sub> ≠ F<sub>2</sub> | sd | 0.07 | 0.00 | 0.10 | 0.10 | 0.14 | 0.08 | 0.11 | 0.17 |
| KS | mean | 2.56 | 99.86 | 32.06 | 67.06 | 73.06 | 60.22 | 89.62 | 73.88 |
| H<sub>0</sub> : F<sub>1</sub> = F<sub>2</sub> H<sub>1</sub> : F<sub>1</sub> ≠ F<sub>2</sub> | sd | 0.05 | 0.01 | 0.15 | 0.14 | 0.26 | 0.16 | 0.09 | 0.25 |
| EB | mean | 4.94 | 0.66 | 5.18 | 87.92 | 0.10 | 78.50 | 87.36 | 96.76 |
| H<sub>0</sub> : E(F<sub>1</sub>) = E(F<sub>2</sub>) H<sub>1</sub> : E(F<sub>1</sub>) ≠ E(F<sub>2</sub>) | sd | 0.07 | 0.02 | 0.07 | 0.10 | 0.01 | 0.13 | 0.11 | 0.07 |
| MWW | mean | 4.68 | 11.22 | 5.76 | 84.96 | 53.36 | 55.50 | 83.94 | 71.36 |
| H<sub>0</sub> : M(F<sub>1</sub>) = M(F<sub>2</sub>) H<sub>1</sub> : M(F<sub>1</sub>) ≠ M(F<sub>2</sub>) | sd | 0.07 | 0.10 | 0.07 | 0.12 | 0.21 | 0.16 | 0.12 | 0.20 |

Furthermore, we investigate the power for cases 2 - 8. KS test and MMD show the relatively good performance such as case 2 where the only distribution shape is different. However, they cannot capture the feature of case 8 where the majority of the two groups are overlapped with each other although 15% of minority of distribution exists. Besides, KS testing fails to detect case 6 where 15% of minority distribution is hyper-methylated. DiffVar shows high power for cases where the variances differ, however, it might capture the other distribution features for the cases with equal variances (case 6), leading to uninterpretable results. EB can appropriately distinguish only the mean difference. MWW can detect case 4, and 7, but cannot detect cases in which the means differ under non-normality. The proposed D^3^M provides preferable results over the eight cases.

On one hand, from this simulation, most of all the distribution features can be captured with D^3^M, which indicates that it works well for the given situations and it can be applied to various cases flexibly. On the other hand, a small sample situation should also be examined. We conduct the simulation considering such a situation and we show the details in supplemental file in S1 and R codes in S2.

## 4 ACTUAL EXAMPLE

### 4.1 Datasets

We apply our method to methylation data of glioblastoma multiforme (GBM) and lower grade glioma (LGG) from The Cancer Genome Atlas (TCGA). GBM is the primary brain tumor that progresses with malignant invasion destroying normal brain tissues (TCGA, 2008), arising through two pathologically distinct routes, *de novo* and as secondary tumors from LGG (Wiencke *et al.*, 2006). In this analysis, we compare the methylation levels in the LGG and GBM groups, and then specify the differential methylation sites. The detection of differential methylation levels is a clue for revealing epigenetic mechanisms of development from LGG to GBM. We focus on mean, variance, and shape differences using EB, DiffVar, KS and D^3^M and compare the results.

Here we briefly describe the datasets and preprocessing as follows. All the samples are hybridized to Illuminas Infinium HumanMethylation450K arrays, including 485,577 CpG sites, which is downloadable from TCGA portal sites. Each CpG site contains 145 samples and 530 samples in GBM and LGG, respectively. We remove CpG sites on the X and Y chromosomes and SNP control probes(rs1–rs65). We also use *HumanMethylation450 v1.2 SNP Update Table* to remove SNP related probes by *minor allele frequency ≤* 1%. As a result, we get 351,932 probes. Missing values in both groups are inferred using R package pcaMethods (Stacklies, 2007) with `pca` functions since DiffVar doesn’t accept missing values and the number of the probes including missing values cannot be ignored. We remove the batch effect using the `ComBat` function in SVA package (Leek, *et al.*, 2012) with default settings. We use batches having more than 2 samples. The details are described in R codes in S3. We finally obtain 141 GBM samples and 530 LGG samples.

### 4.2 Analysis results

Significant differential methylation sites were identified as those having false discovery rate (q-value) (Benjamini, *et al.*, 1995) less than 1%. The Venn diagram (Figure 2 (A)) shows the number of detected probes from the perspective of mean, variance, and shape difference of distributions using EB, DiffVar, KS, and D^3^M, and 279,008, 191,050, 297,493, and 255,317 sites are totally detected, respectively. From Figure 2 (A), most of detected sites with the shape difference are overlapped with the sites by the mean and the variance differences, respectively. However, when focusing on top 1000 significant sites (Figure 2 (B)), the overlaps between them become very small. This suggests that the “signal” of differential methylation sites in terms of *q*-value is quite different from each other. It is important for the further analysis, such as the pathway analysis since we often use the filtered gene set, *e.g.,* using top 1,000 significant sites.

**Fig. 1.**
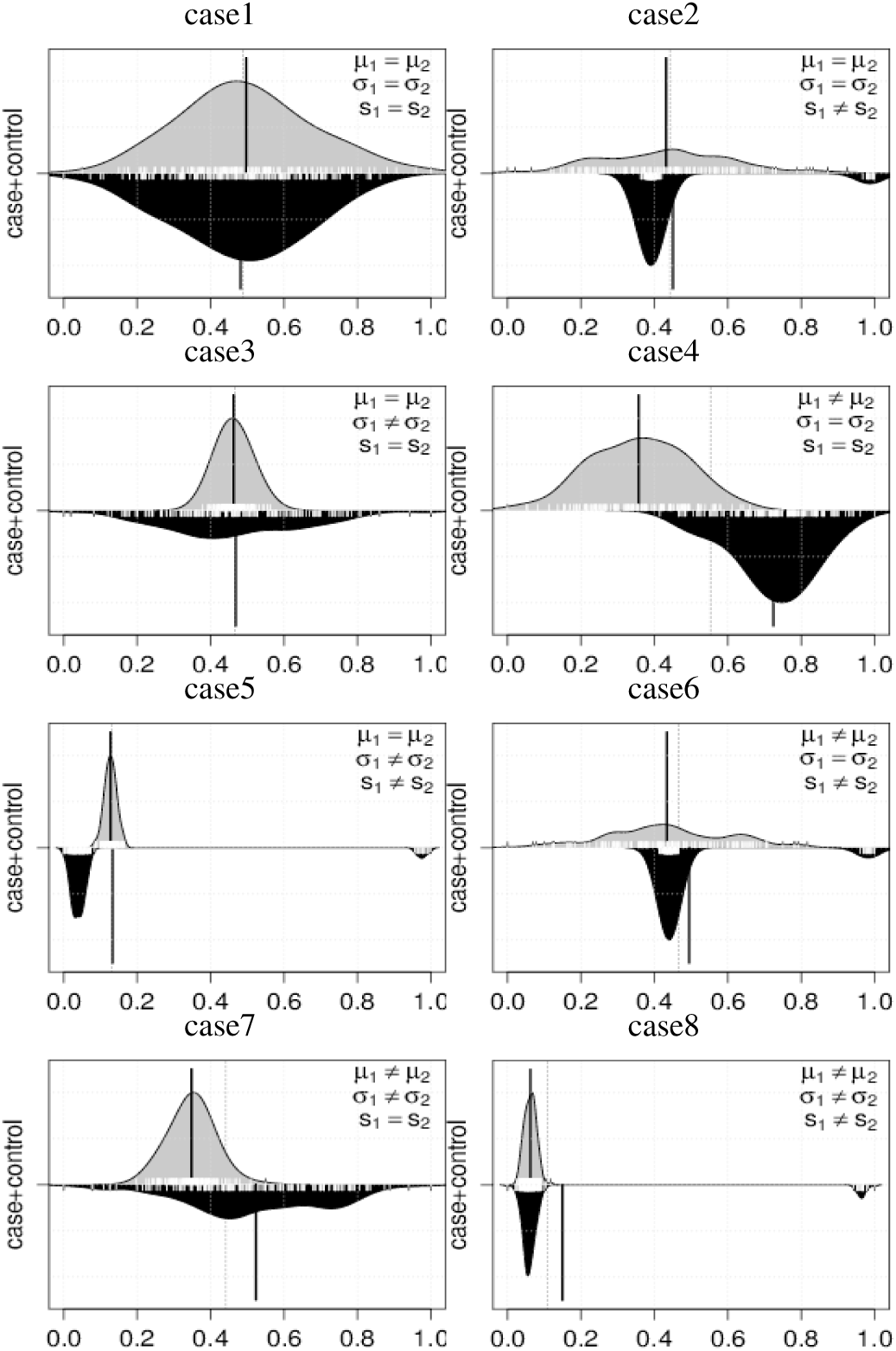
The beanplot of eight cases

**Fig. 2.**
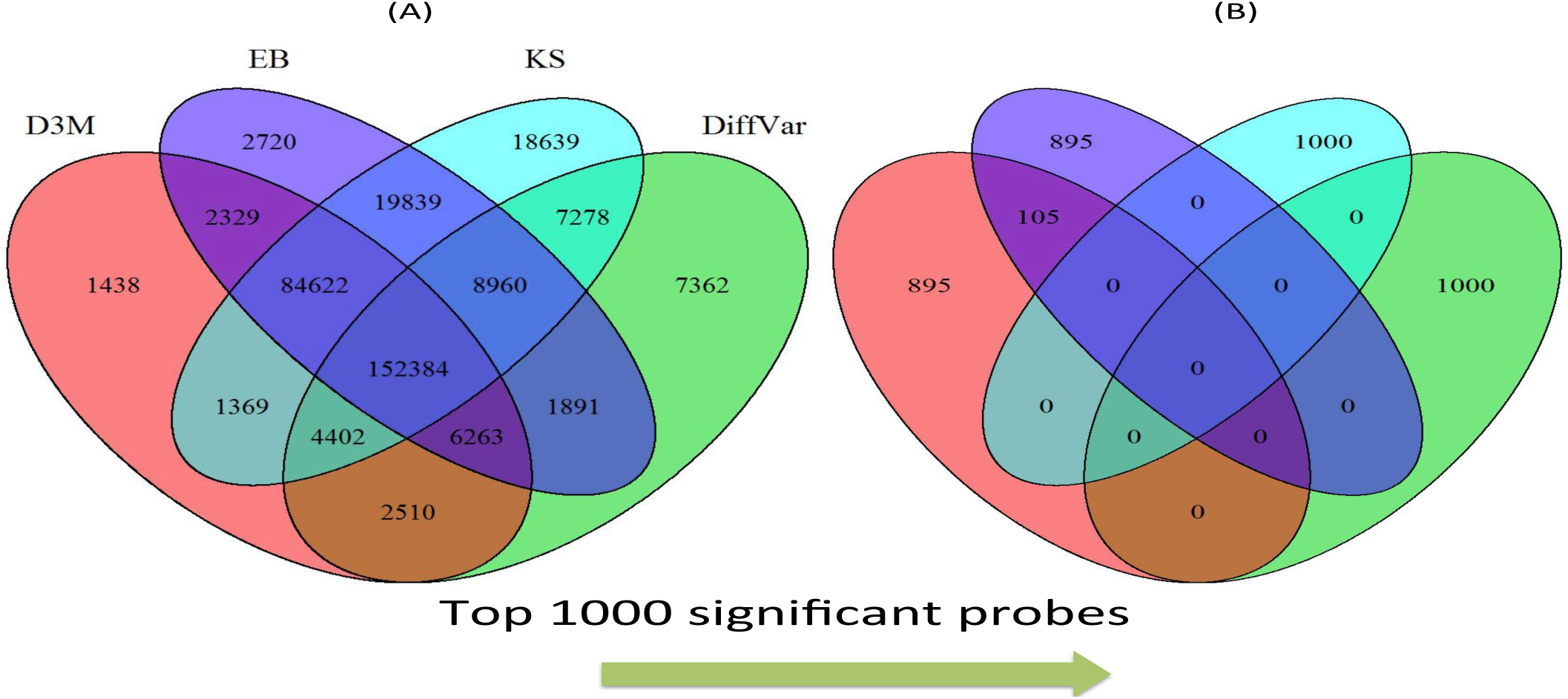
Venn diagram of detected sites at the significance level 1% (A) and of top significant 1000 sites (B).

Among the detected sites, we investigated sites with the smallest 1,000 *q*-value. Heat map and Q–Q plots of the top 1,000 sites are shown in Figures 3 (A) and (B). Comparing heat maps and Q–Q plots, the methylation levels are easy to interpret in the latter. From the Q–Q plot, we could see that the top 1,000 sites tend to be hyper-methylated in LGG (with the reverse in GBM).

**Fig. 3.**
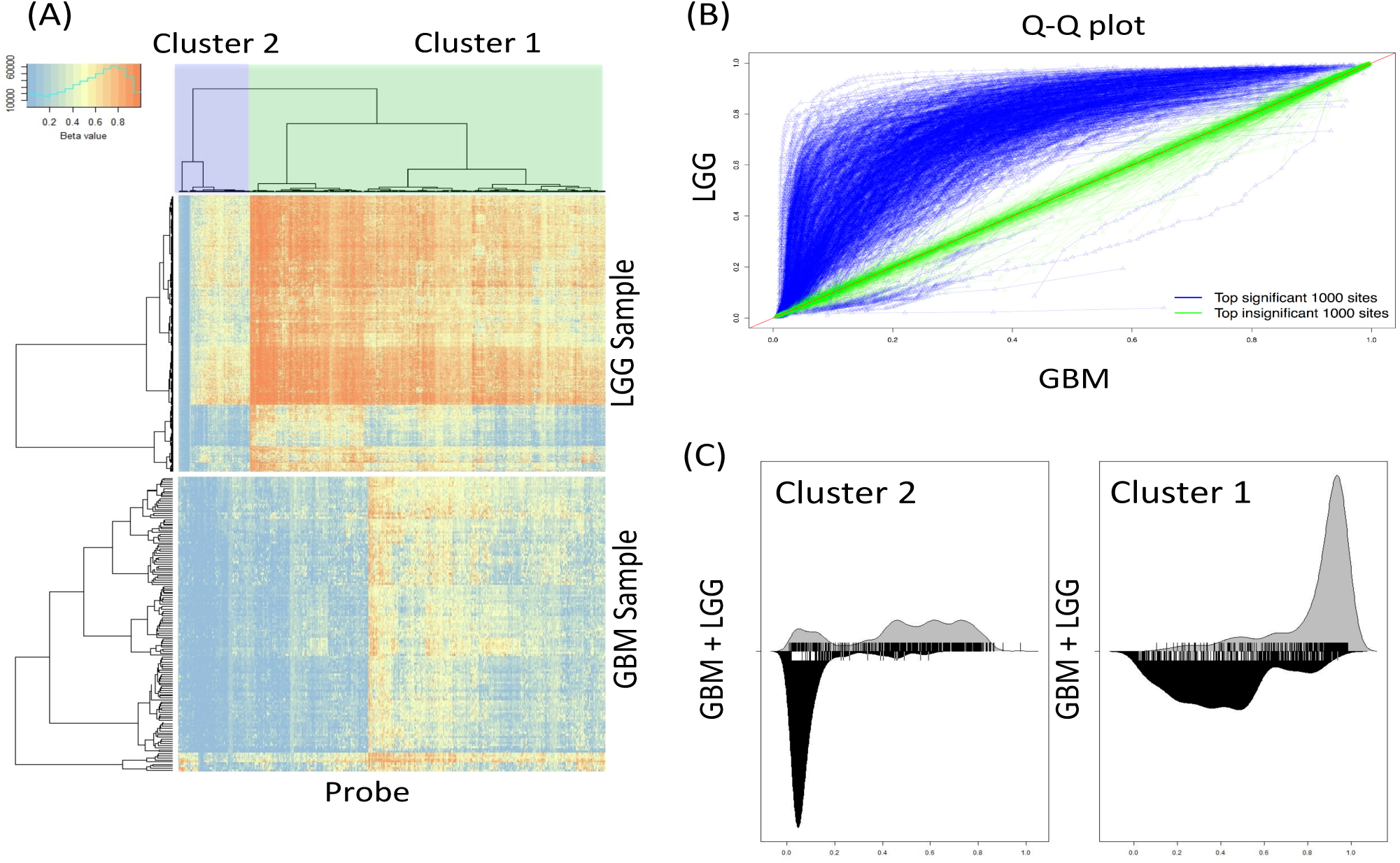
Distributions of the top 1,000 significant probes. (**A**) The heat map of beta values with 141 LGG and 530 GBM samples at each probe respectively. We jointly clustered quantiles of LGG and GBM probes into the two clusters based on the Wasserstein metric (*i.e.,* we clustered curves in Q–Q plots of the top significant probes into two clusters). (**B**) Q–Q plots of beta values between the LGG and GBM probes (blue lines). D^3^M detects the probes with hyper-methylated LGG probes compared with GBM probes. Green lines are the top 1,000 insignificant probes (as negative controls). (**C**) The instances of distributions with Clusters 1 and 2 shown in (A-1) and (A-2), respectively. The distributions appear to be highly heterogeneous especially in LGG probes. D^3^M recognizes strongly heterogeneous distributions and detects such probes as top significant probes.

Among the top significant 1,000 pairs of distributions of GBM and LGG, we can observe that there are mainly two patterns in terms of the Q–Q plot. Then, we cluster the 1000 curves in Q–Q plot into two classes; we consider input data as 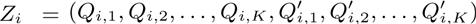 (*i =* 1, 2,…, 1000) and define the dissimilarity as 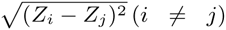 where *Q_i_,_k_* and 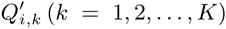 is *k*–th quantile in LGG and GBM group, respectively. Then we apply the standard hierarchical clustering (Clusters 1 and 2 in Figure 3 (A)). Typical distribution examples in each cluster are shown in Figure 3 (C). Clusters 1 and 2 contain 626 and 374 sites, respectively.

Next, we perform enrichment analysis on gene sets in clusters 1 and 2. We used ingenuity pathway analysis (IPA) for 385 and 268 genes in clusters 1 and 2, respectively, and significantly enriched pathways in each cluster using Fisher’s exact test. Table 4 shows five pathways and related genes, ranked with *q*-values in each cluster.

The pathways in clusters 1 and 2 include significant pathways in GBM, which have been previously reported even though we do not include any information on GBM. The axonal guidance signaling pathway in cluster 1 has been suggested as prompting the cell invasion of GBM (Dominique, *et al.*, 2007) and ERK/MAPK Signaling is reported to be up-regulated in GBM (Liu *et al.* 2013). The enrichments of Caveolar-mediated Endocytosis Signaling and Calcium Signaling are studied in (Dong, *et al.*, 2010; Polisetty *et al.*, 2012). The remaining pathways might be explained elsewhere. Our prediction using D^3^M provides a hypothesis that DNA methylation in these pathways might cause the phenotypical difference between GBM and LGG.

We further focus on Wnt in Human Embryonic Stem Cell Pluripotency pathway, and then compare the ranking based on *p*-value by D^3^M with those by other methods. The activation of Wnt family is closely related to cell differentiation of GBM (Rampazzo, *et al.*, 2013). In our analysis, there are 18 Wnt genes on the probes. Among them, six Wnt genes (Wnt2, Wnt2b, Wnt3, Wnt4, Wnt7a, and Wnt 9a) are included in top 1,000 significantly differentially methylated probes using D^3^M.

We investigate the enrichment of the six Wnt genes in top 1,000 significant probes using Fisher’s exact test and we confirm the enrichment of the genes (the details are described in section S-3-2 in supplemental file S1). The six Wnt genes are included in both the clusters 1 and 2, and the majority of the distribution is hypo-methylated and the minority is hyper-methylated in GBM, vice versa in LGG. This suggests that demethylation of Wnt in some LGG might trigger the activation of Wnt family and prompt transformation from LGG to GBM. The ranking of Wnt genes in D^3^M, EB, KS, and DiffVar is shown in Table 3.

**Table 3.** The ranking of the six Wnt gene probes in 351,932 probes. For each method, the upper values are the absolute ranking among 351,932 of the genes, and lower are the their percentages.

|  | Wnt2 | Wnt2b | Wnt3 | Wnt4 | Wnt7a | Wnt9a |
| --- | --- | --- | --- | --- | --- | --- |
| D<sup>3</sup>M | 342<br>(0.10) | 549<br>(0.16) | 591<br>(0.17) | 829<br>(0.24) | 641<br>(0.18) | 887<br>(0.25) |
| EB | 22,541<br>(6.40) | 12,914<br>(3.67) | 3,246<br>(0.92) | 1,811<br>(0.51) | 135<br>(0.04) | 20,747<br>(5.90) |
| KS | 4,754<br>(1.35) | 91,240<br>(25.93) | 50,174<br>(14.26) | 41,589<br>(11.82) | 41,066<br>(11.67) | 84,248<br>(23.94) |
| DiffVar | 56,950<br>(16.18) | 155,621<br>(44.22) | 147,817<br>(42.00) | 146,033<br>(41.49) | 138,928<br>(39.48) | 242,648<br>(68.95) |

**Table 4.**
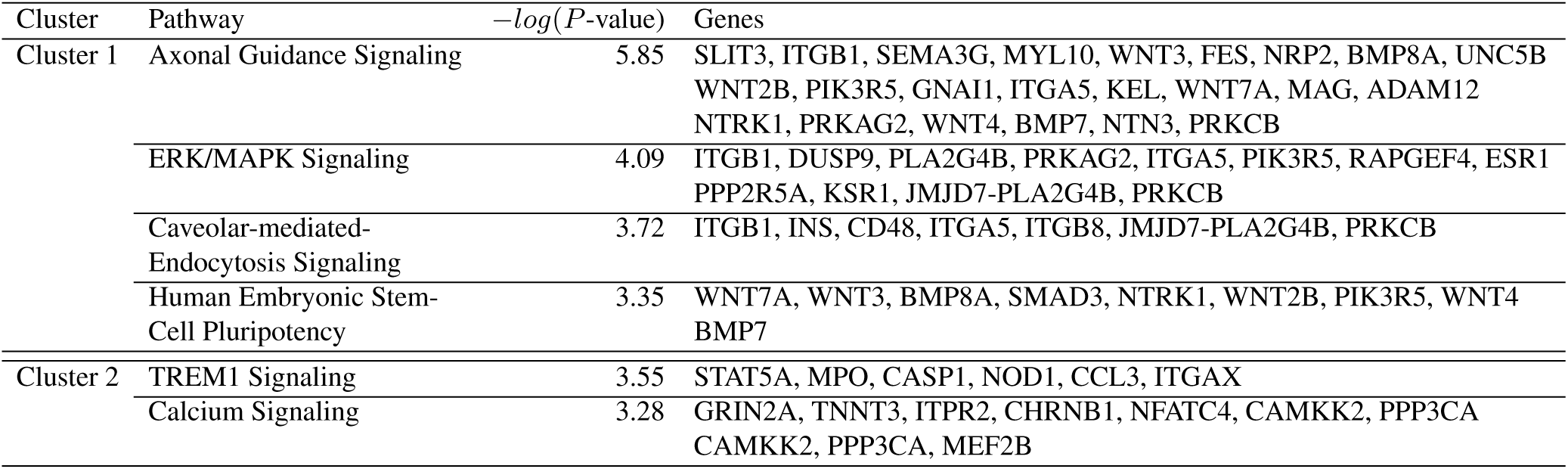
Pathways detected with the proposed method

## 5 DISCUSSION

Here, we summarize the advantages and disadvantages of D^3^M, DiffVar as well as MMD. These methods are designed for detecting differential methylation levels focusing on cancer heterogeneity, which is caused by epigenetic instability and diversity. Cancer heterogeneity can often be confused with outliers. For example, DiffVar fails to detect simulation case 6 as differential methylation, even though we set the variance, but not the mean and the shapes, to be the same for the two groups. This is because DiffVar deals with minority distributions as outliers and evaluates only those in the majority.

In general, the significance of an outlier depends on the context of analysis (Aggarwal, 2013). When an outlier arises from measurement error not relevant to signals of interest, we must remove them prior to analysis. In contrast, when an outlier arises from an unusual event including new findings that we seek, we use them for further analysis. In this case, cancer heterogeneity could be regarded as an abnormal event compared with normal cases, and thus must be included in the analysis.

MMD was originally developed as a distribution-free two sample test based on a kernel. Both the Wasserstein metric and MMD are family of integral probability metrics (Müller, 1997) designed to capture not only such basic difference as those between the means but also such higher-order differences as those between the distribution shapes of the groups. From the simulation in section 3.1, we see that MMD shows good detection performance in cases such as case 2 and 6. D^3^M shows good performance over all eight 8 cases. Specifically, D^3^M shows excellent performance in cases 7 and 8. Considering that case 8 is most frequently observed in actual data, this result is preferable in applications.

In the study of sample size effect in section S-2-1, using the same simulation model as in section 3.1, the result indicates that the superiority of D^3^M to other methods are reduced compared to large sample case, although all the methods decrease in power (Table S.3). On the other hand, MMD retains its power in case 2 and case 6, and so does D^3^M in case 7 and case 8.

We also investigate the power of D^3^M with other simulation models using beta distributions considering small sample size in section S-2-2 and S-2-3 in supplemental file S1. The result indicates that EB and MWW detect differences well in the small samples (*n* = 12, *n* = 20) and so do D^3^M and MMD in the moderate sample sizes (*n* = 28, *n* = 36). In case of unbalanced sample sizes between the case and the control groups (Table S.5), D3M and EB shows the good detection powers.

D^3^M can be applied flexibly to differential methylation problems. Simulation results indicate that D^3^M can detect not only shape differences but also summary statistics differences as effectively as EB and DiffVar, *i.e*., natural results from the decomposition (6). This suggests that if we cannot obtain sufficient power using a simple summary statistics approach, we have other options to add shape information. If we would like to see the results of variance and shape differences simultaneously, we remove the mean from the data using X – E[*X*] in each group at a site before applying D^3^M. This option is provided with R package D^3^M; in the function `D3M::d3m`, the logical parameters of `rm.mean` and `rm.var` exists. If we would like to see the variance and shape differences at the same time, we just set the `rm.mean=T` and `rm.var=F`.

The statistical test of D^3^M relies on resampling and requires computational time to calculate *p*-values. However, we could reduce the resampling time using a semi-parametric approach (Knijnenburg, *et al.*, 2009).

A current limitation of D^3^M is that it deals with univariate distributions. In a case of the study of large sample sizes, we can deal with covariates, such as age and gender, but our method currently does not incorporate them into the model, and the user needs to remove the effects of covariates; for example, using residuals with regression analysis, prior to the analysis. The extension of D^3^M to multivariate distribution relies on the estimation of the Wasserstein metric between the empirical multivariate distributions. The approximation of the Wasserstein metric between the multivariate distributions has been studied recently in (Applegate, *et al.* 2011), and we could derive the null distributions based on that approximation.

D^3^M does not support a spatial correlation in the current form. The spatial information will enhance the power of detection for differential methylated sites. This could be accomplished by spatially weighted average of Wasserstein distance over a fixed range of locus. These extensions of D^3^M will be covered in future work.

## 6 CONCLUSION

In this study, we proposed a novel method, D^3^M, for detecting differential methylation sites based on distribution-valued data. We showed that distribution shape includes interesting information other than that found using mean- and variance-based methods. A simulation study indicated that D^3^M is capable to detect various situations.

In the application to the GBM and LGG dataset in the TCGA cohort, we identified 1,000 sites with the smallest *q*-values. Most of the sites detected by D^3^M show strong heterogeneity and tend to be hyper- and hypo-methylated in LGG and GBM, respectively, as found in previous studies.

Since the GBM and LGG dataset contains a large number of significantly different sites, including 279,008, 191,050, 297,493 and 255,317 sites for D^3^M, EB, KS, and DiffVar, respectively; at the FDR ≤ 1%, it is difficult to understand the methylation levels at these sites. In the future, it would be of interest to develop a method that describes the diversity of methylation levels.

